# A newly identified Duck orbivirus as the etiological agent of egg-production decline in Chinese breeder ducks

**DOI:** 10.1101/2025.10.15.682702

**Authors:** Bing Li, Jianhua Wang, Xuezhi Cui, Dongmin Hao, You Wang, Mingqing Xu, Haixia Chang, Huihui Li, Mingtian Mao, Mian Wu, Chengguang Lu, Jing Tang, Suyun Liang, Zhanbao Guo, Zhengkui Zhou, Youxiang Diao, Shuisheng Hou, Yi Tang

## Abstract

The genus *Orbivirus* comprises double-stranded RNA viruses, many of which are transmitted by arthropods and cause febrile, hemorrhagic, or reproductive diseases in animals. In recent years, unexplained outbreaks characterized by marked egg-production decline have occurred in breeder duck flocks in China. To identify the causative agent, samples were collected from affected farms in multiple regions across China between 2022 and 2025. Representative samples were analyzed by high-throughput sequencing, followed by phylogenetic analysis, transmission electron microscopy, host cell infection assays, and animal challenge experiments.

A previously unreported *Orbivirus*, designated Duck orbivirus (DORV), was identified. Epidemiological investigation indicated that outbreaks mainly affected breeder ducks during the peak laying period (21-55 weeks of age), with higher incidence in summer and autumn. The DORV genome contains ten double-stranded RNA segments encoding seven structural and five nonstructural proteins. Phylogenetic analysis showed that DORV is closely related to Parry’s Lagoon virus (PLV) and Corriparta virus (CORV). Electron microscopy revealed icosahedral particles 30-40 nm in diameter. DORV replicated efficiently in DEF, DEL and C6/36 cells but not in MDCK and LMH cells, suggesting a restricted host range with potential arthropod-borne features. Infection of breeder ducks with DORV-SD01 reduced egg production by 30-40% and caused lesions in the ovary, liver, and spleen.

This study describes a novel duck-origin *Orbivirus* linked to egg-production decline, offering new etiological insights and extending the host spectrum of *Orbivirus*. The results provide a foundation for future surveillance and prevention of emerging waterfowl diseases.

## INTRODUCTION

The genus *Orbivirus* belongs to the family Sedoreoviridae, order Reovirales, class Resentoviricetes, phylum Duplornaviricota, and realm Orthornavirae(Viruses, 2024, Gorman *et al*., 1983). Members of this genus possess a double-stranded RNA (dsRNA) genome composed of ten segments of varying lengths, encoding seven structural proteins (VP1–VP7) and four to five nonstructural proteins (NS1–NS4/5) (Nouda *et al*., 2021, Belaganahalli *et al*., 2015, Roy, 1996, Roy, 1993). The virions exhibit a typical icosahedral symmetry with diameters of approximately 60–80 nm and are non-enveloped, structurally complex RNA viruses (Li *et al*., 2022, Fernández de Castro *et al*., 2014). The genomic segment organization and sequence characteristics are of great importance for phylogenetic analyses within the genus and serve as essential references for molecular epidemiological studies (Kopanke *et al*., 2022).

The genus *Orbivirus* includes multiple viruses of significant veterinary relevance, such as Bluetongue virus (BTV) and African horse sickness virus (AHSV) (Van Rijn *et al*., 2016a, Carpenter *et al*., 2017). These viruses are mainly transmitted by arthropod vectors (e.g., mosquitoes, midges, and ticks) (Belaganahalli *et al*., 2015), infecting both domestic and wild animals and causing diseases characterized by fever, hemorrhage, and arthritis, which pose substantial threats to livestock production and public health. Members of this genus display a wide host range and diverse geographical distributions. Their genomic and antigenic variability increases the complexity of diagnosis and control of diseases (Pascall *et al*., 2020).

In recent years, numerous novel members of the genus *Orbivirus* have been identified. In 2005, Yunnan orbivirus (YUOV) was first isolated from *Culex tritaeniorhynchus* mosquitoes in Yunnan, China, and was found to replicate only in *Aedes albopictus* cells but not in mammalian cell lines(Attoui and H., 2005). In 2007, Middle Point orbivirus (MPOV) was isolated from cattle in Australia(Cowled *et al*., 2007). In 2008, Toggenburg orbivirus (TOV) was detected in goats in Switzerland and identified as a novel serotype of Bluetongue virus(Hofmann *et al*., 2008). In 2011, the genomic features of Umatilla virus (UMAV) were described for the first time(Belaganahalli *et al*., 2011). In 2013, Kapoor et al. reported a new mosquito-borne orbivirus, Sathuvachari virus (SVIV), in Southeast Asia (Kapoor *et al*., 2013). In 2016, Jessica et al. isolated Parry’s Lagoon virus (PLV) from mosquitoes in northern Australia, which showed high genetic and antigenic relatedness to Corriparta virus (CORV) (Harrison *et al*., 2016). Notably, PLV was unable to replicate in various vertebrate cell lines, suggesting a restricted host range. In contrast, CORV, isolated in Namibia in 2021, replicated efficiently in duck-derived AGE1.CR cells, indicating potential cross-host replication capacity (Gonzalez and Knudson, 1987, Guggemos *et al*., 2021). In 2024, Mirolo et al. detected UMAV in the liver tissue of a penguin that died of hepatitis (Mirolo *et al*., 2024), further expanding the known host spectrum of *Orbivirus* (Silva *et al*., 2014).

With the rapid expansion of intensive waterfowl farming, egg-production decline in breeder ducks has become a recurrent and complex clinical issue that severely impacts economic productivity. Since 2022, large-scale outbreaks characterized by a significant drop in egg production have occurred in multiple provinces across China, with the causative agent remaining unknown.

In this study, a previously unrecognized orbivirus was identified and isolated from the ovarian tissue of breeder ducks exhibiting reduced egg production. High-throughput sequencing, virus isolation, and systematic analyses confirmed that this pathogen represents a novel member of the genus *Orbivirus*, designated Duck orbivirus (DORV). Comprehensive genomic sequencing, phylogenetic analysis, transmission electron microscopy, and multi-host cell culture assays revealed the genomic structure, evolutionary relationships, and replication characteristics of DORV, suggesting its potential for arthropod-borne transmission. Animal infection experiments demonstrated that infection with DORV-SD01 caused a pronounced decline in egg production and typical pathological lesions in the ovary, liver, and spleen of breeder ducks.

In conclusion, this study provides the first systematic characterization of the molecular and biological properties of a novel duck-origin orbivirus and elucidates its pathogenic mechanisms. These findings establish DORV as a key etiological agent of egg-production decline in breeder ducks and provide a theoretical foundation for targeted prevention strategies and the surveillance of emerging viral diseases in waterfowl.

## MATERIALS AND METHODS

### Sample collection and processing

Since 2022, clinical samples have been collected from breeder duck farms in multiple regions of China, including Shandong, Jiangsu, Henan, Guangxi, and Inner Mongolia. Affected flocks showed reduced egg production, increased mortality, or slow recovery of laying rate. Some ducks also had poor feed intake, soft or small eggs, mild respiratory signs, tearing, diarrhea, and occasional joint swelling. Liver, spleen, oviduct, ovarian follicle, and intestinal tissues were collected and stored at −80 °C. One representative ovarian sample from a Shandong flock in 2022 was used for genome analysis, virus isolation, and pathogenicity test.

### RNA extraction and quality control

Total RNA was extracted from the supernatants of tissue homogenates using an automated nucleic acid extraction system (NanoMigBio-96, China) and a magnetic bead-based viral RNA extraction kit (NMG1811, China) according to the manufacturer’s instructions. RNA concentration and purity (A260/280 and A260/230) were measured using a NanoDrop One spectrophotometer (Thermo Fisher Scientific, USA), and RNA integrity was assessed using a Bioanalyzer 2100 (Agilent Technologies, USA) with an RNA 6000 Nano Kit. Samples with an RNA integrity number (RIN) ≥ 7 were considered suitable for library preparation, and 3 µg of total RNA was used as input.

### Library construction and sequencing

RNA-seq libraries were constructed using the NEBNext Ultra^TM^ Directional RNA Library Prep Kit for Illumina (New England Biolabs, USA). Ribosomal RNA was removed prior to library preparation, and first- and second-strand cDNA synthesis was performed using random primers. The average insert size (approximately 250–300 bp) was verified using an Agilent Bioanalyzer 2100 (Agilent Technologies, USA). Paired-end sequencing (2 × 150 bp) was performed on the Illumina NovaSeq 6000 platform.

### Quality control, read processing, and assembly

Raw reads were assessed for quality using FastQC (v0.11.9). Adapter sequences and low-quality bases (Phred score < 20) were trimmed using Trimmomatic (v0.39), and reads shorter than 50 bp were discarded to obtain high-quality clean reads. De novo assembly was performed using CLC Genomics Workbench (v21.0.6; QIAGEN Digital Insights, Germany) to generate contigs. Putative viral contigs were identified and annotated by BLASTn search against the NCBI Virus (nt subset) database. The resulting genome was classified as complete or near-complete based on sequence coverage and alignment depth.

### Virus detection

Total RNA was extracted, and complementary DNA (cDNA) was synthesized using a premixed reverse transcription kit (AG11728, China). Conventional PCR amplification was performed with specific primers (Table S1) using the 2× PCR Master Mix (CW3009M, China) according to the manufacturer’s instructions. Amplified products were analyzed by 1.5–2.0% agarose gel electrophoresis to detect the target fragments. Quantitative real-time PCR was performed on a LightCycler 96 system (Roche, Switzerland) using specific primers (Table S1) and the premixed qPCR Master Mix (CW3360M, China) following the manufacturer’s protocol.

### Viral similarity analysis

The viral genome sequence obtained in this study was compared with representative reference sequences of related species and strains available in GenBank (see Table S2). Multiple sequence alignment was performed using ClustalW implemented in MegAlign (DNASTAR, USA). Misaligned and gap-rich regions were manually inspected and trimmed. Pairwise nucleotide identities were calculated from the curated alignments, and a nucleotide similarity matrix was generated. The results were visualized in R (v4.5.0) using the pheatmap package, with hierarchical clustering based on Euclidean distance and the complete-linkage algorithm.

### Phylogenetic analysis

Multiple sequence alignment was performed using ClustalW implemented in MEGA 11, and phylogenetic trees were constructed using the neighbor-joining (NJ) method with 1,000 bootstrap replicates to evaluate the robustness of tree topology.

### Virus isolation and cell culture

Tissue homogenates were centrifuged at 8,000 × g for 10 min, and the supernatants were filtered through a 0.22-µm membrane filter (Merck Millipore, USA). The filtrates were inoculated onto duck embryo fibroblast (DEF) and duck embryo hepatocyte (DEH) monolayers cultured in DMEM/F12 medium (Gibco, USA) supplemented with 10% fetal bovine serum (FBS; Gibco, USA) and 1% penicillin–streptomycin (Gibco, USA). After 1 h of adsorption, the inoculum was replaced with maintenance medium containing 2% FBS, and the cultures were incubated at 37 °C with 5% CO_2_. Cells were examined daily for cytopathic effects (CPE), and supernatants were collected 1–5 days post-inoculation (dpi). If no apparent CPE was observed, the cultures were blindly passaged up to three times.

### Transmission electron microscopy

The purified viral supernatant was centrifuged at 8,000 × g for 10 min to remove impurities, and the clarified supernatant was adsorbed onto carbon-coated copper grids (200 mesh) for 5 min. The grids were negatively stained with 2% phosphotungstic acid (pH 7.0) for 1 min, blotted dry with filter paper, and air-dried. Samples were observed and photographed using a transmission electron microscope (Hitachi TEM system, Japan).

### Viral growth kinetics assay

Duck embryo fibroblast (DEF), duck embryo hepatocyte (DEH), mosquito-derived C6/36, chicken-derived LMH, and canine MDCK cells were seeded at a density of 1 × 10^6^ cells/mL in 24-well plates and cultured until approximately 80% confluence. The cells were then infected with DORV-SD01 at a multiplicity of infection (MOI) of 1, incubated for 1 h for virus adsorption, and the inoculum was replaced with maintenance medium. At 1, 2, 3, 4, and 5 days post-infection (dpi), both the cells and supernatants were collected. Viral RNA was extracted and quantified by qPCR, and the viral copy numbers at each time point were calculated. The viral growth curves were plotted with the log_10_ (copies/mL) values on the y-axis.

### Mosquito sample collection and processing

Mosquitoes were captured using hand nets in both DORV-affected farms and geographically distant residential areas (n = 3, with 10 mosquitoes pooled per sample). After morphological identification, the mosquito samples were stored at −80 °C until use for DORV detection by qPCR.

### Pathogenicity experiment

A total of sixty 22-week-old healthy breeder ducks at peak laying were randomly divided into two groups (30 ducks per group). The experimental group was inoculated via the wing vein with 0.5 mL of viral suspension (1 × 10^6^ RNA copies/mL), while the control group received an equal volume of sterile saline. The experiment lasted 30 days, during which ducks were monitored daily for clinical signs, feed and water intake, and egg production. All procedures were approved by the Institutional Animal Care and Use Committee of the Institute of Animal Sciences, Chinese Academy of Agricultural Sciences (approval no. IAS2025-129).

### Histopathological examination

When the infected group exhibited a significant decline in egg production, ducks were randomly selected from both groups and humanely euthanized under isoflurane anesthesia by cervical dislocation. The ovaries, oviducts, liver, and spleen were collected and examined for gross pathological changes. Tissues were fixed in 10% neutral-buffered formalin, embedded in paraffin, sectioned at 3–5 μm, and stained with hematoxylin and eosin (H&E). Histopathological changes were observed and photographed using a light microscope (Nikon, Japan).

## RESULTS

### Pathogenic characteristics

Since 2022, a group disease characterized by a marked decline in egg production has emerged in several breeder duck farms in Shandong and Inner Mongolia. Affected flocks were mainly adult breeder ducks at the peak of egg laying. Within 1–2 weeks after onset, egg production dropped sharply by 30–40% (Figure 1D). Ducks in affected flocks showed depression, reduced feed intake, and increased proportions of soft-shelled and small eggs. The disease lasted approximately 2–3 weeks, and while some flocks partially recovered, the egg production level generally did not return to normal. A mild increase in mortality was also observed.

**Figure 1.**
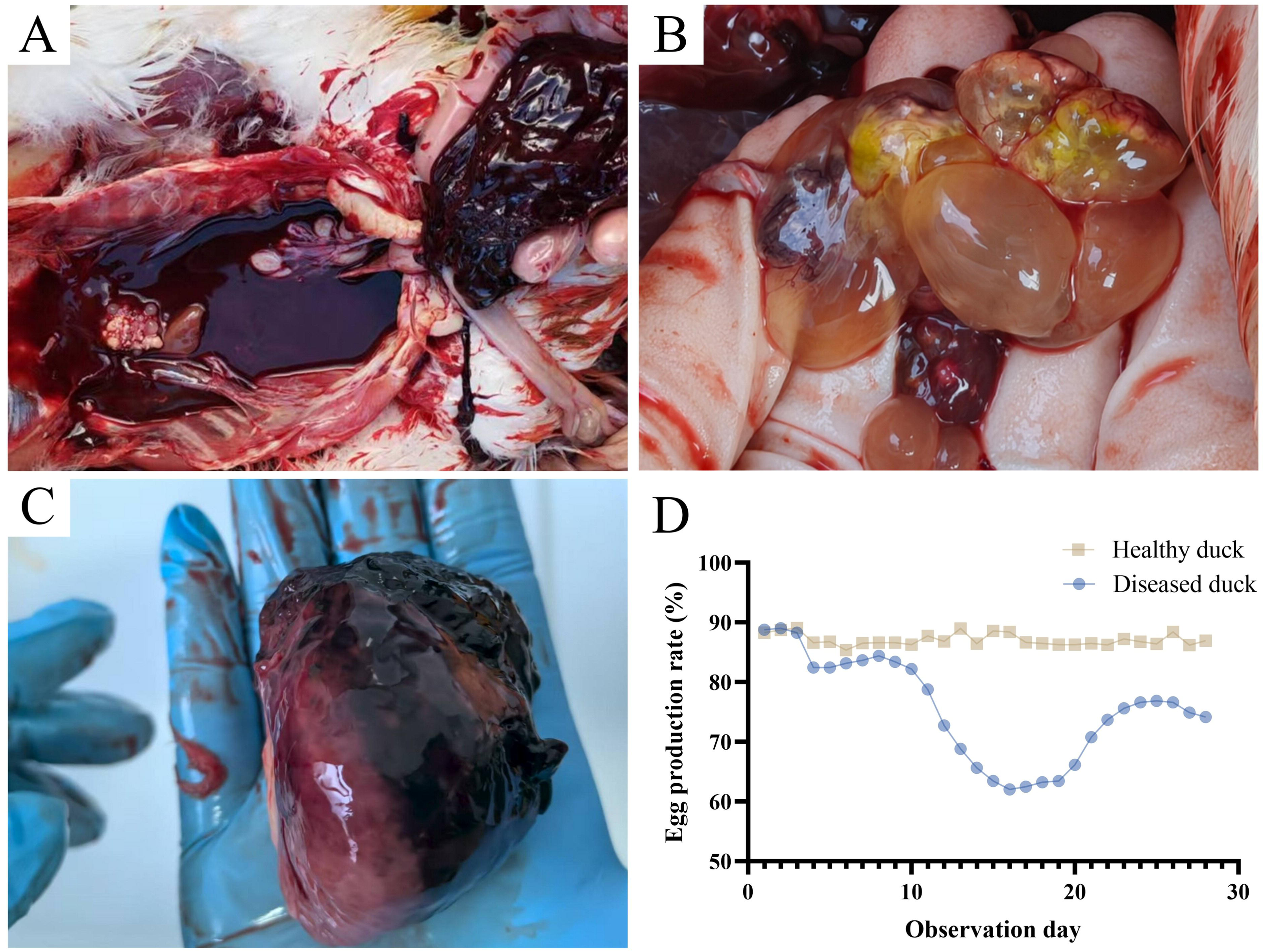
Representative pathological features of affected breeder ducks. (A) Abdominal cavity filled with blood; liver showing hemorrhage, enlargement, and friability; (B) Ovarian follicles showing edema, liquefaction, necrosis, and hemorrhage, accompanied by caseous necrotic material; (C) Enlarged and congested spleen with firm texture; (D) Comparison of egg production rates between healthy and affected ducks (data from Ningyang, Shandong).

Gross examination revealed that the most prominent lesions occurred in the liver, spleen, and reproductive system (Figure 1A-C). Severe hemorrhage in the abdominal cavity was a characteristic lesion distinguishing this disease. The liver was enlarged, friable, and darkened, with multifocal or patchy hemorrhage. The spleen was markedly swollen and firm, sometimes ruptured with a mottled appearance. The number of ovarian follicles was significantly reduced; some were edematous, congested, necrotic, or hemorrhagic, often containing caseous necrotic material, and in severe cases, follicular liquefaction was observed. The oviduct was hyperemic and edematous, with yellow or gray-white caseous exudate in the lumen. Some individuals also exhibited systemic lesions, including ascites, myocardial hemorrhage, pulmonary edema, and necrotizing enteritis.

### Virus identification and epidemiological analysis

The representative clinical sample collected from Shandong Province in 2022 was subjected to second-generation genome sequencing. Sequence alignment against the NCBI viral nucleotide (nt) database revealed that the genome shared the highest similarity with viruses of the genus *Orbivirus*, including Parry Lagoon virus (PLV), Corriparta virus (CORV), and Acado virus (ACAV). All alignments showed an E-value of 0, indicating an extremely low probability of random matches and a high degree of sequence similarity. The pathogen responsible for the decline in egg production in breeder ducks was identified as a novel *Orbivirus* species with ducks as the natural host, designated as Duck orbivirus (DORV).

Since 2022, a total of 32 clinical samples showing decreased egg production have been collected from various regions of China. All samples tested positive for DORV by both PCR and qPCR assays. Spatially, the cases were mainly distributed in the major duck-breeding areas of China, with 53.1% from Shandong Province and 25.0% from the Inner Mongolia Autonomous Region (Figure 2A). Temporally, the number of suspected cases increased steadily from the first detection in 2022, reaching a peak in 2025, when “isolated egg production decline” became the predominant clinical presentation (Figure 2B). Most affected ducks were breeding ducks aged 21–55 weeks (accounting for 75% of all cases), corresponding to the early-to-late laying period (Figure 2C). DORV infections occurred throughout the year, with the highest incidence in summer and autumn (40.6% and 34.4%, respectively), and lower rates in winter and spring (15.6% and 9.4%). Overall, most cases occurred between June and November (75.0%) (Figure 2E).

**Figure 2.**
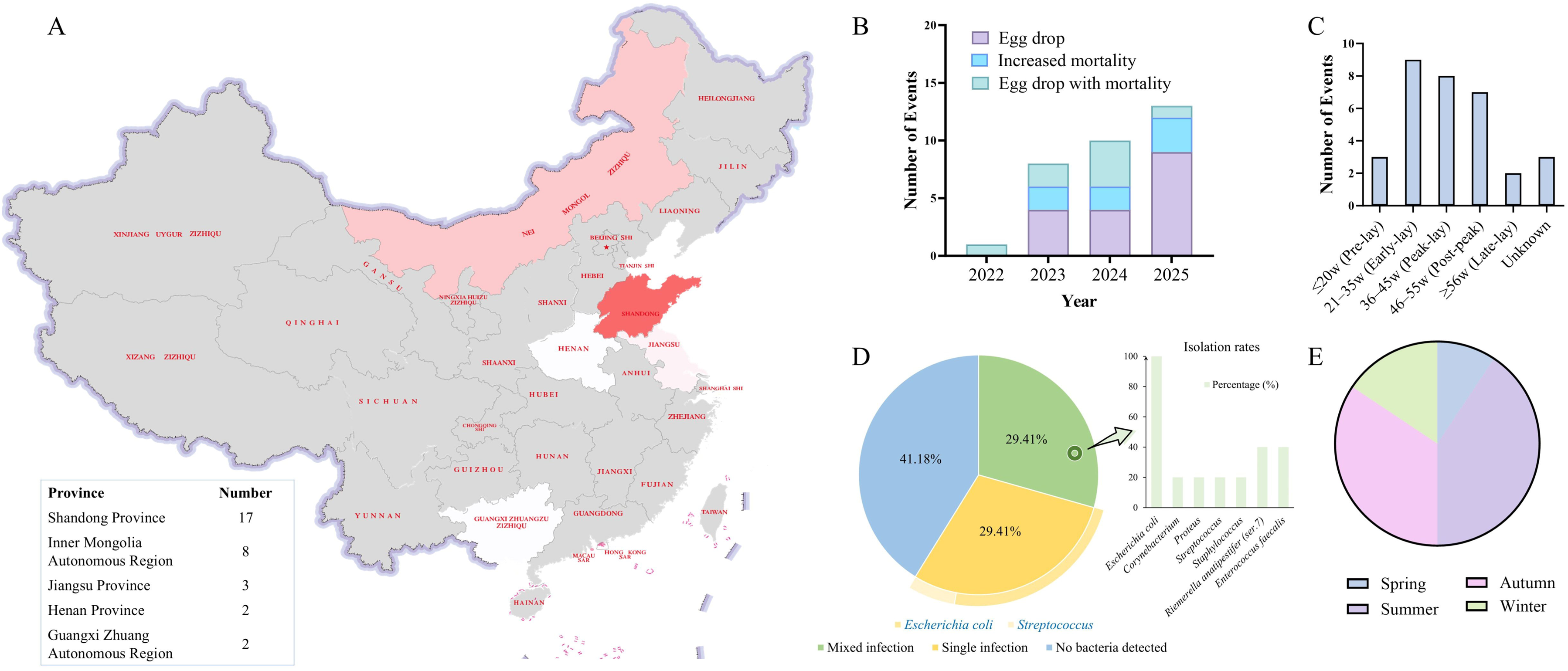
Geographic and epidemiological characteristics of DORV infection in breeder ducks. (A) Geographic distribution of DORV-positive cases in China; (B) Annual distribution of positive events from 2022 to 2025; (C) Distribution of cases by age group; (D) Bacterial infections detected in DORV-positive samples; (E) Seasonal distribution of cases, with the corresponding months for spring, summer, autumn, and winter being months 3-5, 6-8, 9-11, and 12-2, respectively.

**Figure 3.**
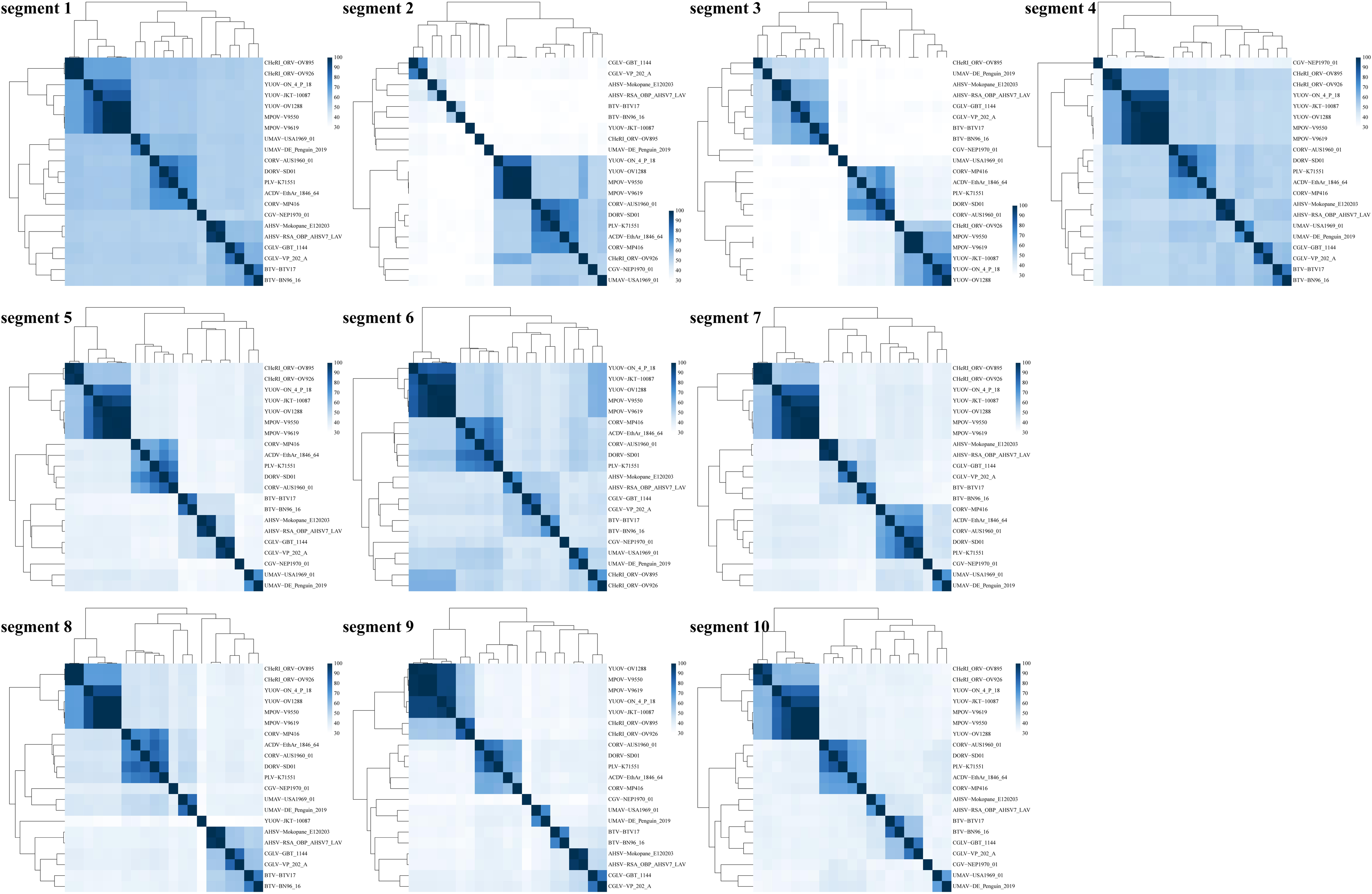
Heatmap of nucleotide sequence similarity between DORV and reference *Orbivirus* strains. Each subpanel represents one of the ten double-stranded RNA genome segments of DORV (Segments 1–10). The color intensity indicates the level of nucleotide sequence similarity among different virus strains, with darker blue denoting higher similarity. The main diagonal indicates self-comparison.

Pathogenetic analysis revealed that Approximately 30% of samples had mixed bacterial infections, 30% had single infections, and 40% were negative for bacteria. Among the isolated bacterial strains, Escherichia coli was the most common, accounting for 80% of single infections and being almost universally present in mixed infections (Figure 2D).

### Genome characteristics analysis

The genome of Duck orbivirus (DORV) consists of 10 segments of double-stranded RNA, ranging in length from 717 bp (segment 10) to 3,873 bp (segment 1). Except for segments 9 and 10, each segment contains a single major open reading frame (ORF); segments 9 and 10 each encode two proteins. In total, 12 viral proteins were identified, including seven structural proteins (VP1–VP7) and five nonstructural proteins (NS1, NS2, NS3, NS3a, and NS4), with predicted sizes ranging from 153 to 1,291 amino acids. The encoded proteins include RNA polymerase, core proteins, and capsid proteins, which are involved in replication and transcription, structural assembly, and regulatory functions (Table 1).

### Genomic similarity analysis

Comparative analysis based on sequence similarity and phylogenetic clustering of the 10 genomic segments of DORV showed that the virus exhibited a stable clustering pattern across most segments, forming a distinct phylogenetic lineage. Within the genus *Orbivirus*, DORV exhibited the highest sequence identity with PLV, CORV, and ACDV (Figure 2).

DORV showed the highest similarity to PLV-K71551 in seven of the 10 genomic segments (VP1: 89.956%, VP2: 91.333%, VP4: 87.975%, NS1: 85.342%, NS2: 92.294%, VP6/NS4: 90.725%, and NS3/NS3a: 88.028%) (Table 2). The highest similarity in VP3 and VP5 was observed with CORV-AUS1960_01 (82.768% and 82.544%, respectively), while VP7 showed the highest similarity to ACDV-EthAr_1846_64 (85.032%).

Compared with other animal-derived *Orbiviruses*, including BTV, AHSV, Changuinola virus (CGV), Chuzan virus (CHUV), and YUOV, DORV showed substantial sequence divergence among all genomic segments. The nucleotide identity ranged from 29.9% to 52.5%, with the lowest identity (29.9%) observed in segment 2 (VP2) (Table 3).

Overall, DORV displayed typical genomic features of *Orbivirus*, but exhibited significant divergence from all known reference strains across every segment, supporting its classification as a novel duck-origin *Orbivirus* species that is evolutionarily closest to the PLV lineage.

### Genetic Evolutionary Analysis

Comparative phylogenetic analyses were conducted for the 10 genomic segments (Segment 1–10) of DORV-SD01 (Figure 4). The results showed that most phylogenetic trees exhibited strong bootstrap support for deep-level branching, whereas the shallow-level relationships among DORV, PLV, CORV, and ACDV showed lower support values. The main uncertainty was concentrated in determining the closest phylogenetic affinity of DORV. Noticeable differences in scale bars and branch lengths among segments indicated distinct evolutionary rates. The short internal branches within the DORV-PLV/CORV/ACDV clusters suggested close genetic relatedness and relatively recent divergence, whereas their long-distance separation from AHSV, BTV, CGLV, and CGV reflected distant evolutionary divergence.

**Figure 4.**
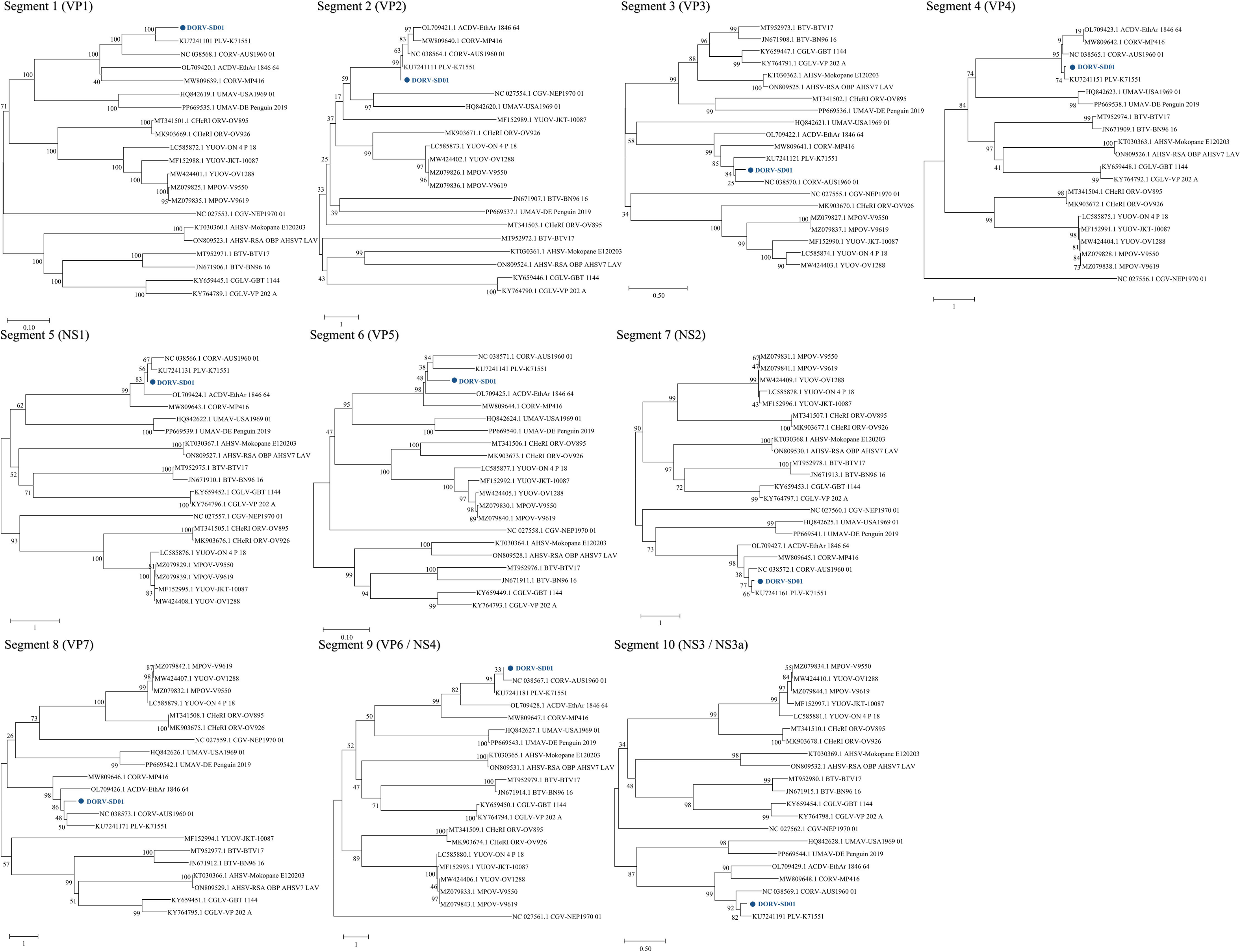
Phylogenetic analysis of the 10 genomic segments of DORV-SD01. Blue circles indicate DORV-SD01. Bootstrap values are shown at branch nodes, and the scale bar represents the number of nucleotide substitutions per site.

In most segments, DORV-SD01 clustered tightly with PLV (K71551), including VP1, VP2, VP4, NS1, NS2, VP6/NS4, and NS3/NS3a. Among these, VP2 and VP6/NS4 showed short proximal branches with relatively low support values, suggesting minor uncertainty in the fine-scale topology within the cluster. In Segment 3, DORV-SD01 formed a sister relationship with CORV (AUS1960_01), while Segments 5 and 7 grouped DORV-SD01 with ACDV (EthAr_1846-64), with bootstrap support ranging from moderate to high. Collectively, DORV represents a novel duck-origin *Orbivirus* species that is most closely related to the PLV lineage but exhibits significant genomic divergence.

### Morphological and Growth Characteristics Analysis

The viral strain isolated from samples collected in Shandong Province in 2022 was designated as DORV-SD01. Negative-staining transmission electron microscopy revealed that DORV-SD01 virions were uniform, spherical particles with clearly defined edges and homogeneous electron density, measuring approximately 30–40 nm in diameter (Figure 5B), consistent with the typical morphology of the genus *Orbivirus*. To evaluate host cell adaptability, DORV-SD01 was inoculated into culture systems derived from different species for replication assays. The results showed that DORV-SD01 infection caused marked cytopathic effects (CPE) in duck embryo fibroblast (DEF) cells and duck embryo liver (DEL) cells, including cellular shrinkage, detachment, deformation, loose intercellular connections, disordered arrangement, and reduced density. In mosquito C6/36 cells, massive detachment and prominent vacuolation were observed. In contrast, no apparent CPE was observed in chicken hepatocellular carcinoma (LMH) cells or canine kidney (MDCK) cells. Infected duck embryos exhibited growth retardation, surface congestion, darkened coloration, and hemorrhagic foci on the abdominal and thoracic regions, with multiple subcutaneous petechiae, particularly large dark red ecchymoses around the abdomen and liver (Figure 5A).

**Figure 5.**
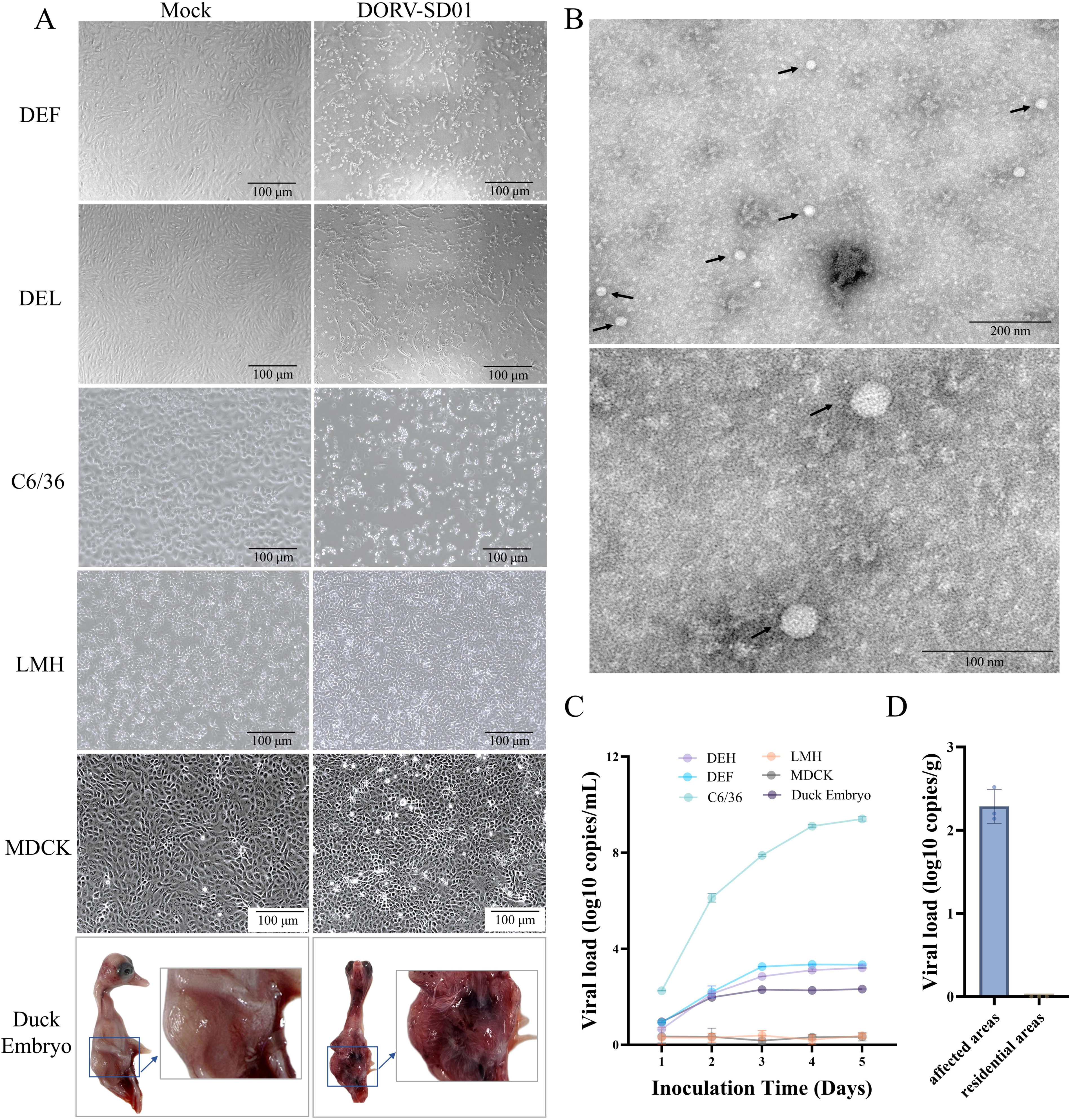
Morphological characteristics and replication capacity of DORV-SD01 in different hosts. (A) Cytopathic effects and pathogenic manifestations in different cells and duck embryos infected with DORV-SD01; (B) Negative-staining transmission electron microscopy of DORV-SD01; arrows indicate viral particles; Enlarged view showing clearly defined particle boundaries and uniform electron density; (C) Growth curves of DORV-SD01 in different cells and duck embryos. Viral loads were determined by qPCR and expressed as log_10_ (copies/mL); (D) Comparison of DORV viral loads in mosquito samples collected from different regions. Error bars represent standard deviations (n = 3).

Quantitative PCR was used to assess viral replication levels in different cells and duck embryos (Figure 5C). The results showed that DORV-SD01 replicated most efficiently in C6/36 cells, with viral loads increasing rapidly from 1 day post-infection (dpi) and peaking at 5 dpi (approximately 10^9.4^ copies/mL), indicating robust replication in mosquito cells. Viral amplification was also detected in DEF and DEL cells, rising sharply at 2–3 dpi and then stabilizing. Viral loads in duck embryos were slightly lower than in DEF and DEL cells but remained consistently detectable. In contrast, viral replication in LMH and MDCK cells was extremely limited, with no significant amplification observed.

Furthermore, DORV detection in mosquito samples (*Culex* spp.) collected from DORV-affected and geographically distant residential areas revealed that viral RNA was detectable at low concentrations only in mosquitoes from affected regions (Fig. 5D).

### Pathogenicity

The clinical manifestations, egg production dynamics, and major tissue lesions of breeder ducks infected with DORV-SD01 were systematically monitored from 1 to 28 days post infection (dpi) (Figure 6). Results showed that the egg production rate of the infected group was comparable to that of the PBS control group during the early stage of infection (1–2 dpi, approximately 86%–90%). However, egg production began to decline markedly from 3 dpi and reached its lowest level at 8–12 dpi (approximately 53%–56%), representing a reduction of about 35 percentage points compared with the control. Although production gradually recovered thereafter, it remained significantly lower than the control (approximately 90%) at 28 dpi. DORV-SD01 infection caused a persistent decrease in egg-laying performance.

**Figure 6.**
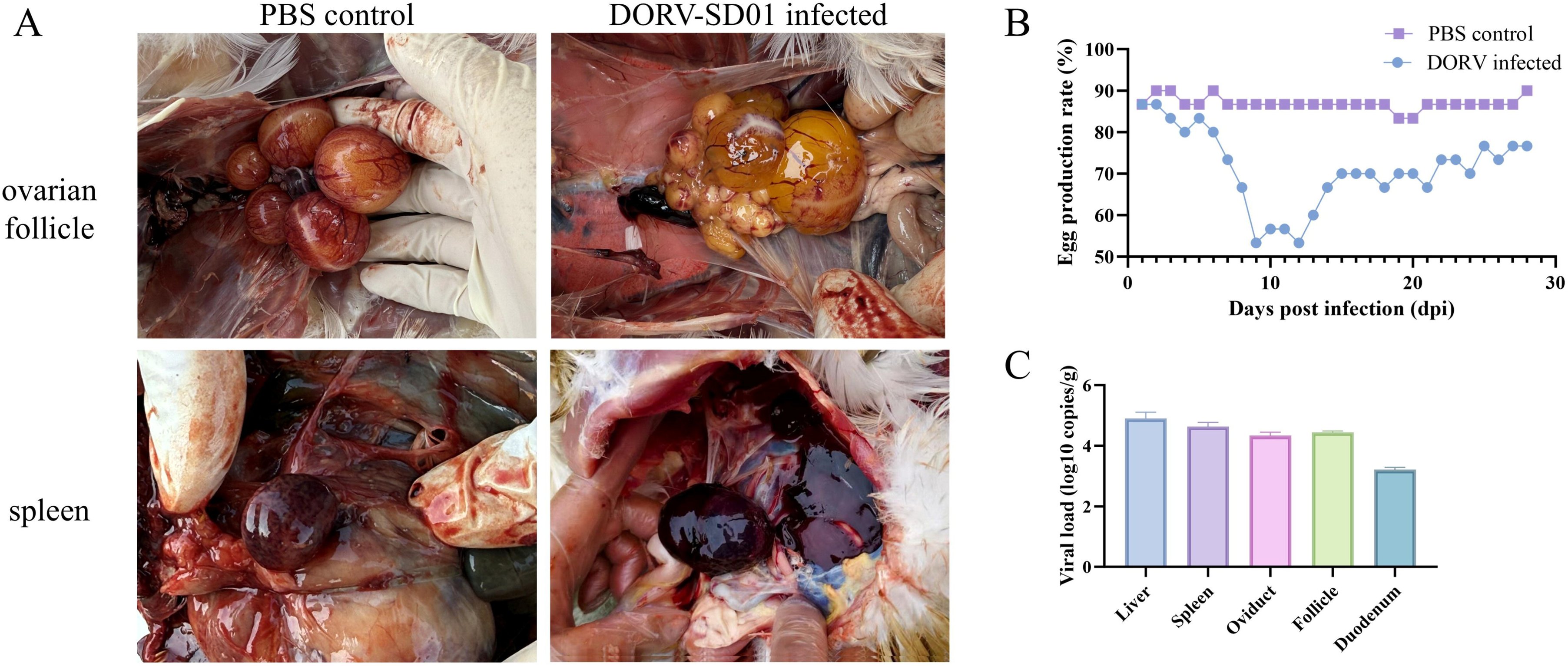
Pathogenicity of DORV-SD01 infection in breeder ducks. (A) Representative lesions in major tissues after infection. (B) Changes in egg production rate following infection. (C) Viral load distribution across different tissues.

Necropsy revealed that the ovaries of infected ducks were swollen, with multiple follicles of varying sizes distributed on the surface. Some follicles appeared malformed with indistinct contours, rupture, and yellow coagulated exudates. The ovarian tissue was fragile and loosely structured, with indistinct boundaries from surrounding tissues, often accompanied by hemorrhage and necrosis. The spleen was markedly enlarged with diffuse hemorrhage, and the liver exhibited severe hemorrhagic lesions. Quantitative detection showed that viral loads were high in multiple tissues, with the liver, spleen, oviduct, and ovarian follicles containing the highest levels, followed by the duodenum. DORV-SD01 infection caused reproductive dysfunction, follicular necrosis, and splenic hemorrhage, leading to a significant reduction in egg production performance.

### Histopathological characteristics

Histopathological examination revealed that DORV infection caused typical lesions in multiple organs, including the ovary, liver, and spleen (Figure 7).

**Figure 7.**
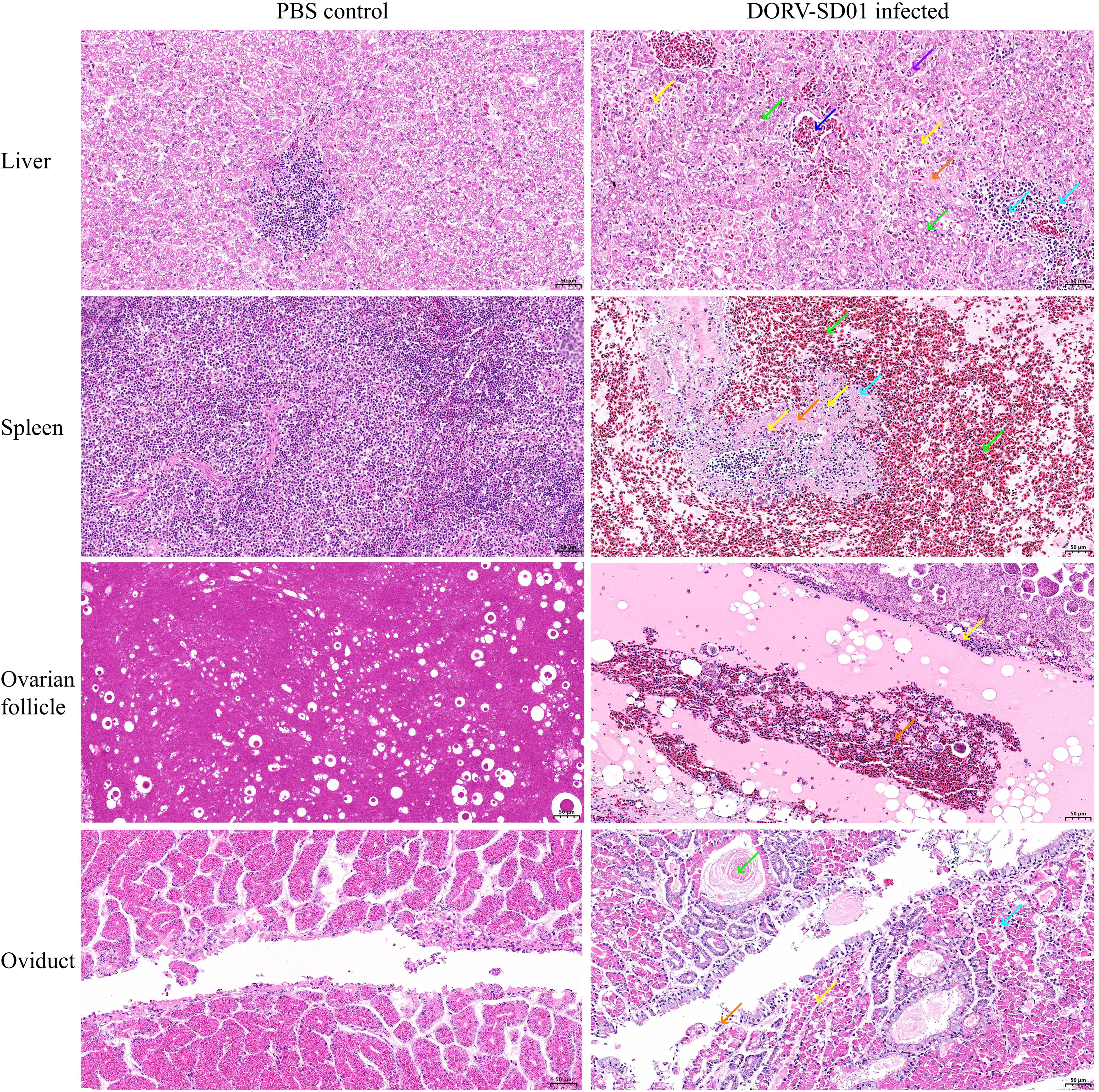
Histopathological changes in breeding ducks infected with DORV-SD01. Scale bar = 50 μm.

Liver: In the control group, hepatic capsules were intact, and hepatocytes and hepatic sinusoids were radially arranged around the central vein, with only mild fatty degeneration. In the infected group, hepatic architecture was severely disrupted, and extensive eosinophilic exudation was observed within the hepatic sinusoids. Focal lymphocytic infiltration was frequently detected around the central vein and portal areas. Hepatocytes were irregularly arranged, with numerous focal necrotic cells exhibiting nuclear pyknosis and fragmentation. Some hepatocytes showed vacuolar degeneration, with small round vacuoles in the cytoplasm. Pronounced fatty degeneration was also noted in some cells, characterized by variably sized lipid vacuoles within the cytoplasm. A small number of hepatocytes were edematous, with cell swelling and pale, loose cytoplasm. Multiple foci of lymphocytic infiltration were evident.

Spleen: In the control group, splenic architecture was intact, with no apparent abnormalities. In the infected group, splenic tissue exhibited extensive structural disorganization and widespread hemorrhage, with a marked reduction in splenic cells. Numerous necrotic cells with nuclear fragmentation were observed, accompanied by eosinophilic exudation and infiltration of granulocytes.

Ovarian follicle: In the control group, ovarian follicles were abundant, structurally intact, and morphologically normal. In the infected group, the number of follicles was significantly reduced, and follicular structures were indistinct. The follicles appeared homogeneously eosinophilic, with increased staining intensity, and focal fibroplasia was observed. A large amount of yolk material accumulated within the follicular cavity, surrounded by a homogeneous, acellular yolk membrane. Local necrotic cellular debris and multiple foci of hemorrhage were present within the yolk membrane, accompanied by scattered erythrocytes.

Oviduct: In the control group, oviductal epithelial cells were regularly arranged with clear structural organization. In the infected group, numerous mucosal epithelial cells exhibited edema, swelling, and pale, loose cytoplasm. The glands in the lamina propria were irregularly arranged, with poorly defined structures, and the glandular epithelial cell nuclei were shrunken and deeply stained. The muscle layer was thinner, and necrotic cellular debris was observed.

## DISCUSSION

Since 2022, breeder duck flocks in Shandong and Inner Mongolia, China, have repeatedly exhibited cases characterized by a marked decline in egg production. Routine diagnostic testing failed to detect common duck pathogens such as avian influenza virus, duck plague virus, or duck Tembusu virus, suggesting the presence of a previously unrecognized etiological agent. High-throughput sequencing combined with comparison against the NCBI viral nucleotide (nt) database revealed high sequence similarity to viruses of the genus *Orbivirus*. This newly identified pathogen was designated Duck orbivirus (DORV), and the isolated strain was named DORV-SD01.

Epidemiologically, PCR and qPCR testing showed that all 32 case samples with similar clinical features collected from multiple regions in China were DORV-positive. Spatially, positive samples were mainly concentrated in two core breeder-duck regions, Shandong (53.1%) and the Inner Mongolia Autonomous Region (25.0%), with additional detections in Jiangsu and Henan, indicating multifocal occurrence and regional clustering across major production areas. Since the first detection in 2022, case numbers have increased annually, with a seasonal peak in summer and autumn (June to November). These patterns suggest that temperature, humidity, and diurnal temperature variation may influence viral activity and host susceptibility. Approximately 60% of positive samples showed bacterial co-infections, most commonly *Escherichia coli*, implying potential DORV-associated immunosuppression and that secondary bacterial infections may exacerbate tissue inflammation and reduce laying performance. Therefore, on-farm control strategies should address both viral and bacterial factors.

Genomic analysis showed that DORV comprises 10 segments of double-stranded RNA encoding 12 structural and nonstructural proteins, consistent with canonical *Orbivirus* genome architecture. Among reference *Orbivirus* strains, DORV exhibited the highest sequence identity with PLV, CORV, and ACDV. Seven segments showed the highest identity to PLV-K71551 (up to 91.3%), whereas VP3, VP5, and VP7 displayed marked divergence, indicating a distinct evolutionary branch with host-adaptation features. Such divergence may alter tissue tropism or receptor engagement, enabling efficient replication in the avian reproductive tract (Sanjuán and Domingo-Calap, 2016). Overall, DORV is a novel waterfowl-origin *Orbivirus* closely related to PLV yet genetically distinct.

Morphologically, DORV particles are spherical, measuring approximately 30–40 nm in diameter, which is smaller than the typical 60–80 nm reported for *Orbivirus*. DORV replicated efficiently in mosquito C6/36 cells, duck DEF and DEL cells, and duck embryos, with the most rapid replication observed in C6/36 cells. In contrast, replication was minimal or undetectable in chicken LMH and canine MDCK cells, suggesting strong adaptation to both mosquito and duck cells. Although interclass replication has not been observed, environmental adaptation and genetic variation may eventually expand its host range (Barrett and Schluter, 2008), emphasizing the importance of continued cross-host surveillance and vector control.

Considering the geographic distribution and seasonal patterns of DORV, together with the well-known arthropod-borne nature of the *Orbivirus* genus (MacLachlan and Guthrie, 2010), DORV is most likely a mosquito-borne virus. In this study, DORV RNA was detected in mosquito samples collected from affected regions, suggesting that *Culex* spp. may be involved in its natural transmission cycle. Moreover, the virus replicated efficiently in C6/36 cells derived from *Aedes albopictus*, indicating that *Aedes* mosquitoes may also serve as potential vectors. These findings suggest that DORV may exhibit a multi-mosquito-species transmission pattern, although further experimental evidence is required to confirm this hypothesis. More than 30 *Orbivirus* species have been described, infecting ruminants (Omoga *et al*., 2023), equids (Tirosh-Levy and Steinman, 2022), camels (Mohd Jaafar *et al*., 2014), birds (Peňazziová *et al*., 2022), and rodents (Feng *et al*., 2025, Fagre *et al*., 2019). BTV, AHSV, and epizootic hemorrhagic disease virus (EHDV) are biologically transmitted by biting midges (*Culicoides*), with replication in the vector and dissemination to salivary glands (Maan *et al*., 2020, Mellor, 1990). Several avian-origin *Orbivirus* strains have been isolated from mosquitoes: YUOV from *Culex tritaeniorhynchus* (Attoui and H., 2005); UMAV from *Culex pipiens* and *Culex tarsalis* (Belaganahalli *et al*., 2011); PLV and Stretch Lagoon orbivirus (SLOV) repeatedly from *Culex annulirostris* (Belaganahalli *et al*., 2011, Harrison *et al*., 2016); and similar reports exist for CORV. Koyama Hill virus (KHV), isolated in Japan, also shows high titers in mosquito cells with restricted replication in mammalian cells (Ejiri *et al*., 2014), consistent with the growth phenotype of DORV. Although some *Orbivirus* species, such as BTV, can also spread through non-vector routes including direct contact, transplacental transmission, or oral exposure, these are considered exceptional events. DORV, however, appears to possess population-level transmissibility, as evidenced by clustered outbreaks observed among breeder duck flocks (Figure 8) (Weaver and Barrett, 2004).

**Figure 8.**
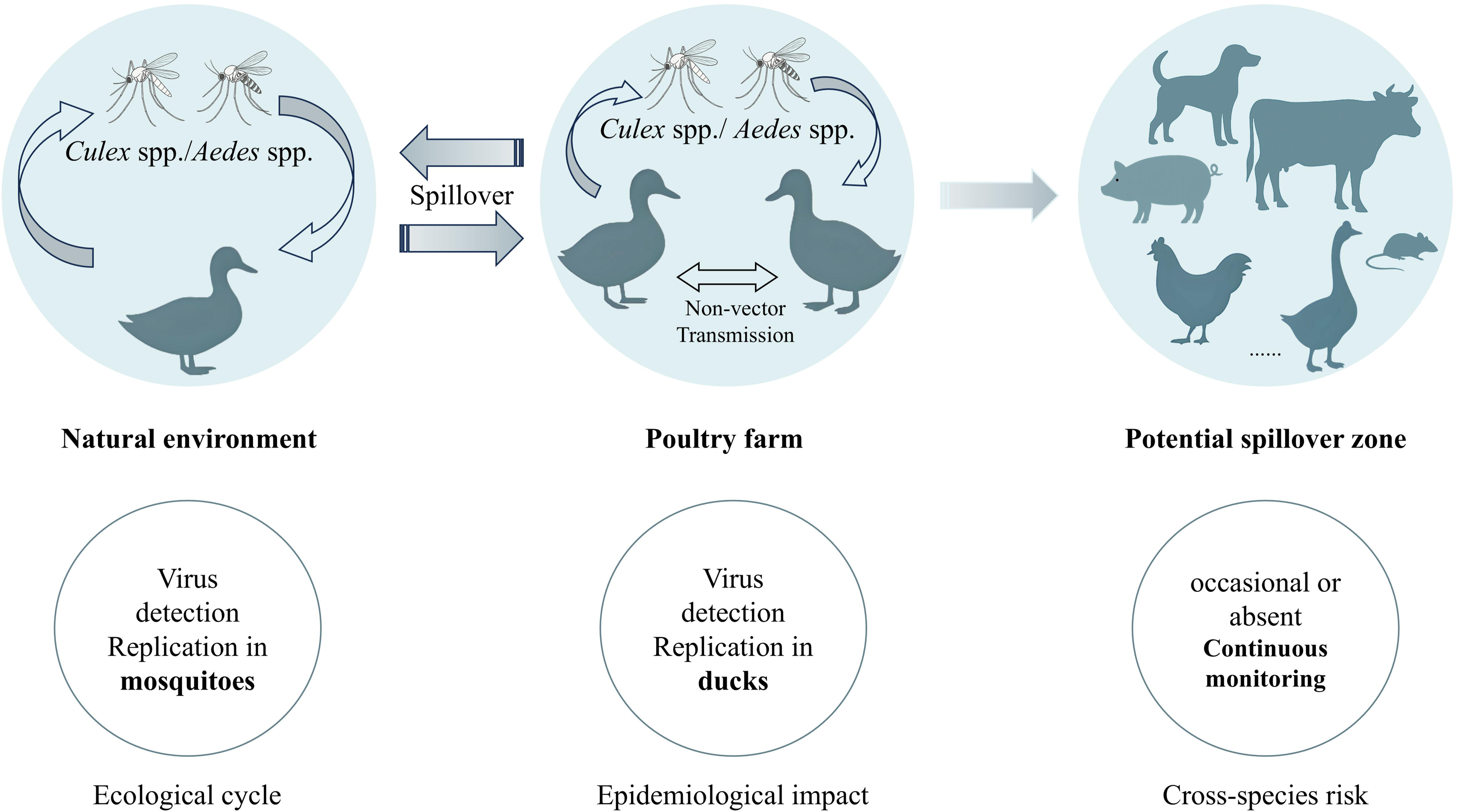
Hypothesized multi-scale transmission model of DORV. The model illustrates potential ecological and epidemiological transmission cycles of DORV among mosquitoes (*Culex spp.*), ducks, and other susceptible hosts. Arrows indicate possible transmission routes, including vector-mediated and non-vector transmission. The rightmost panel highlights potential cross-species spillover zones that require continuous monitoring.

At the pathogenic level, DORV infection is characterized by persistent reductions in egg production and ovarian necrosis. Gross pathology includes ovarian swelling, follicular rupture with caseous necrosis, pronounced hyperemia and edema of the oviduct, and marked hepatic and splenic hemorrhage. Viral-load profiling indicated the highest concentrations in liver, spleen, and reproductive tissues; thus, ovary, oviduct, liver, and spleen are recommended as priority tissues for clinical sampling. The reproductive tropism of DORV may involve direct tissue injury and virus-induced immune stress or endocrine dysregulation, causing impaired folliculogenesis and ovulatory dysfunction, and ultimately leading to substantial egg-production losses and inferior egg quality. Given that BTV can cause embryonic death, abortion, and fetal malformations in ruminants, and EHDV can undergo vertical transmission in vectors and affect midge reproduction (Van Rijn *et al*., 2016b, Maclachlan *et al*., 2019), DORV may also possess vertical transmission potential that warrants targeted investigation.

Histopathology further delineated the injury pattern: in the liver, extensive necrosis and steatosis were accompanied by eosinophilic exudation within sinusoids and inflammatory infiltration; in the spleen, architectural disruption with widespread hemorrhage and marked cellular depletion was evident; in the ovary, follicle numbers were reduced with heightened eosinophilia and focal hemorrhage; in the oviduct, epithelial edema, glandular disorganization, and necrotic debris were present. These lesions indicate not only localized tissue necrosis but also systemic vasculitis and immunosuppression. Splenic damage may disrupt B-cell zones and antibody production, limiting the development of immunological memory and predisposing to secondary bacterial infection (Vale and Schroeder Jr, 2010). The coexistence of eosinophilic infiltration and necrotic debris in the ovary and oviduct suggests that viral infection may induce activation of the necroptosis pathway (Pendl and Schmidt, 2024).

In summary, DORV is a newly identified duck-origin *Orbivirus* that exhibits distinct evolutionary characteristics in its molecular profile, host adaptation, and pathogenic spectrum. DORV targets the reproductive system, leading to follicular necrosis and persistent impairment of egg production. Given its high replication capacity in mosquito cells and the detection of positive mosquito samples in affected regions, DORV is presumed to be maintained through a waterfowl–mosquito–waterfowl transmission cycle and to spread within high-density farming environments. Future studies should focus on elucidating both vector and non-vector transmission mechanisms and determining whether vertical transmission occurs; evaluating host susceptibility and carrier spectra across different species; identifying viral replication sites in the ovary and liver and clarifying their interactions with host immune and endocrine responses; and establishing sensitive surveillance systems to assess the epidemiological dynamics and economic impact of DORV in China.

In conclusion, this study is the first to identify and comprehensively characterize a novel duck-origin *Orbivirus* with reproductive system–targeted pathogenicity. The virus exhibits marked tissue tropism, systemic pathological effects, and potential vector-borne transmission within its host, suggesting that DORV may represent an emerging and significant viral pathogen in the waterfowl industry, warranting further research on prevention strategies and molecular mechanisms.

## Acknowledgement

The authors gratefully acknowledge all collaborators who contributed to this work. We sincerely thank Shandong Hekangyuan Biological Breeding Co., Ltd. (Jinan, China) for providing breeder ducks and technical support.

## Ethics statement

All animal experiments were conducted in accordance with the *Guidelines for Experimental Animals* issued by the Ministry of Science and Technology of China and designed to minimize animal suffering. Euthanasia was performed by exposure to carbon dioxide (CO_2_) to ensure humane treatment. Reporting followed the ARRIVE guidelines. Ethical approval for animal use was granted by the Experimental Animal Welfare and Ethics Committee of the Institute of Animal Sciences, Chinese Academy of Agricultural Sciences (approval no. IAS2025-129).

---

**Table 1. Genome organization and functional annotation of DORV segments.**

**Table 2. Top three sequence identities between DORV genomic segments and reference *Orbivirus* strains.**

**Table 3. Lowest three sequence identities between DORV genomic segments and reference *Orbivirus* strains.**

