## Supplemental Data 1 for "A newly identified Duck orbivirus as the etiological agent of egg-production decline in Chinese breeder ducks"

**Table 1. Genome organization and functional annotation of DORV segments.**

| Segment | Segment length (bp) | ORF length (bp) | ORF position (bp) | Encoded protein | Predicted protein length (aa) | Protein function |
| --- | --- | --- | --- | --- | --- | --- |
| 1 | 3873 | 3873 | 1–3873 | VP1 | 1291 | RNA-dependent RNA polymerase |
| 2 | 2850 | 2850 | 1–2850 | VP2 | 950 | Major outer capsid attachment protein; determinant of antigenic variation and serotype specificity |
| 3 | 2224 | 2193 | 21–2213 | VP3 | 731 | Core scaffold protein (T2); encapsidates genome and participates in core assembly |
| 4 | 1997 | 1917 | 13–1929 | VP4 | 638 | mRNA capping enzyme with guanylyltransferase and methyltransferase activities (cap1 formation) |
| 5 | 1916 | 1770 | 12–1781 | NS1 | 590 | Nonstructural protein; forms tubules and regulates viral protein translation and assembly |
| 6 | 1659 | 1584 | 45–1628 | VP5 | 528 | Membrane-penetration protein; mediates endosomal escape and membrane permeabilization |
| 7 | 1140 | 1065 | 6–1070 | VP7 | 355 | Core-surface layer protein (T=13 lattice); stabilizes the core |
| 8 | 1126 | 1116 | 11–1126 | NS2 | 372 | Major component of viral inclusion bodies (VIBs); binds RNA and recruits core proteins |
| 9 | 1100 | 1035 | 23–1057 | VP6 | 345 | RNA-dependent ATPase/helicase |
|  |  | 459 | 117–575 | NS4 | 153 | Nonstructural protein 4; involved in modulation of host response |
| 10 | 717 | 717 | 1–717 | NS3 | 239 | Membrane-associated glycoprotein with two transmembrane domains; mediates viral budding and egress. |

**Table 2. Top three sequence identities between DORV genomic segments and reference *Orbivirus* strains.**

| **Segment** | **Highest 1** | | **Highest 2** | | **Highest 3** | |
| --- | --- | --- | --- | --- | --- | --- |
|  | **Strain** | **%Identity** | **Strain** | **%Identity** | **Strain** | **%Identity** |
| Segment 1 (VP1) | PLV-K71551 | 89.956 | CORV-AUS1960_01 | 77.66 | ACDV-EthAr_1846_64 | 72.201 |
| Segment 2 (VP2) | PLV-K71551 | 91.333 | CORV-AUS1960_01 | 80.175 | ACDV-EthAr_1846_64 | 75.368 |
| Segment 3 (VP3) | CORV-AUS1960_01 | 82.768 | PLV-K71551 | 81.592 | ACDV-EthAr_1846_64 | 73.813 |
| Segment 4 (VP4) | PLV-K71551 | 87.975 | CORV-AUS1960_01 | 78.813 | ACDV-EthAr_1846_64 | 71.042 |
| Segment 5 (NS1) | PLV-K71551 | 85.342 | CORV-AUS1960_01 | 85.273 | ACDV-EthAr_1846_64 | 77.044 |
| Segment 6 (VP5) | CORV-AUS1960_01 | 82.544 | ACDV-EthAr_1846_64 | 82.294 | PLV-K71551 | 82.197 |
| Segment 7 (NS2) | PLV-K71551 | 92.294 | CORV-AUS1960_01 | 80.373 | ACDV-EthAr_1846_64 | 71.492 |
| Segment 8 (VP7) | ACDV-EthAr_1846_64 | 85.032 | PLV-K71551 | 84.868 | CORV-AUS1960_01 | 82.883 |
| Segment 9 (VP6 / NS4) | PLV-K71551 | 90.725 | CORV-AUS1960_01 | 80.134 | ACDV-EthAr_1846_64 | 65.385 |
| Segment 10 (NS3 / NS3a) | PLV-K71551 | 88.028 | CORV-AUS1960_01 | 80.986 | ACDV-EthAr_1846_64 | 69.859 |

**Table 3. Lowest three sequence identities between DORV genomic segments and reference *Orbivirus* strains.**

| **Segment** | **Lowest 1** | | **Lowest 2** | | **Lowest 3** | |
| --- | --- | --- | --- | --- | --- | --- |
|  | **Strain** | **%Identity** | **Strain** | **%Identity** | **Strain** | **%Identity** |
| Segment 1 (VP1) | AHSV-RSA_OBP_AHSV7_LAV | 52.418 | CGV-NEP1970_01 | 52.447 | AHSV-Mokopane_E120203 | 52.522 |
| Segment 2 (VP2) | AHSV-Mokopane_E120203 | 29.885 | BTV-BN96_16 | 30.985 | YUOV-JKT-10087 | 31.377 |
| Segment 3 (VP3) | CGV-NEP1970_01 | 30.848 | BTV-BN96_16 | 31.071 | AHSV-RSA_OBP_AHSV7_LAV | 31.116 |
| Segment 4 (VP4) | CGV-NEP1970_01 | 36.385 | AHSV-RSA_OBP_AHSV7_LAV | 50.873 | YUOV-ON_4_P_18 | 51.182 |
| Segment 5 (NS1) | BTV-BN96_16 | 35.633 | CGLV-GBT_1144 | 35.907 | CGLV-VP_202_A | 36.353 |
| Segment 6 (VP5) | BTV-BTV17 | 44.578 | AHSV-RSA_OBP_AHSV7_LAV | 45.49 | AHSV-Mokopane_E120203 | 45.556 |
| Segment 7 (NS2) | CGLV-VP_202_A | 36.814 | BTV-BN96_16 | 37.068 | BTV-BTV17 | 37.628 |
| Segment 8 (VP7) | YUOV-JKT-10087 | 35.054 | BTV-BN96_16 | 38.659 | BTV-BTV17 | 38.751 |
| Segment 9 (VP6 / NS4) | CGV-NEP1970_01 | 32.394 | MPOV-V9550 | 34.917 | MPOV-V9619 | 34.917 |
| Segment 10 (NS3 / NS3a) | CHeRI_ORV-OV895 | 36.656 | AHSV-Mokopane_E120203 | 38.146 | CHeRI_ORV-OV926 | 38.344 |
