## Supplemental Data 2 for "A newly identified Duck orbivirus as the etiological agent of egg-production decline in Chinese breeder ducks"

Table S1. Primers used for PCR and RT-qPCR detection of Duck orbivirus (DORV).

| Target | Primer name | Sequence (5′→3′) | Product size (bp) | Application |
| --- | --- | --- | --- | --- |
| DORV | DORV-F | GATACTGGAAGAGATGCTACA | 565 | PCR |
|  | DORV-R | CTGACGCTTGAGACACATA |  |  |
| DORV | DORV-q-F | GATGGAAGTAGCGGTTCTC | 115 | qPCR |
|  | DORV-q-R | GTCAGTCTGTTCACGAATATC |  |  |

Table S2. Information of reference sequences involved in genomic comparison.

| Virus (abbr.) | Strain/Isolate | Accessions (Segments 1–10) |
| --- | --- | --- |
| Duck orbivirus (DORV) | SD01 | PX398457-PX398466 |
| Acado virus (ACDV) | EthAr 1846-64 | OL709420-OL709429 |
| African horse sickness virus (AHSV) | Mokopane-E120203 | KT030360-KT030369 |
| African horse sickness virus (AHSV) | RSA/OBP/AHSV7/LAV | ON809523-ON809532 |
| Bluetongue virus (BTV) | BTV17 | MT952971-MT952980 |
| Bluetongue virus (BTV) | BN96/16 | JN671906-JN671915 |
| CHeRI orbivirus (CHeRI-ORV) | OV895 | MT341501-MT341510 |
| CHeRI orbivirus (CHeRI-ORV) | OV926 | MK903669-MK903678 |
| Changuinola virus (CGLV) | GBT 1144 | KY659445-KY659454 |
| Changuinola virus (CGLV) | VP 202 A | KY764789-KY764798 |
| Chobar Gorge virus (CGV) | NEP1970/01 | NC_027553-NC_027562 |
| Corriparta virus (CORV) | MP416-NA-2018 | MW809639-MW809648 |
| Corriparta virus (CORV) | AUS1960/01 | NC_038564-NC_038573 |
| Middle Point orbivirus (MPOV) | V9550 | MZ079825-MZ079834 |
| Middle Point orbivirus (MPOV) | V9619 | MZ079835-MZ079844 |
| Parry's Lagoon virus (PLV) | K71551 | KU724110-KU724119 |
| Umatilla virus (UMAV) | USA1969/01 | HQ842619-HQ842628 |
| Umatilla virus (UMAV) | DE/Penguin/2019 | PP669535-PP669544 |
| Yunnan orbivirus (YUOV) | ON-4/P/18 | LC585872-LC585881 |
| Yunnan orbivirus (YUOV) | JKT-10087 | MF152988-MF152997 |
| Yunnan orbivirus (YUOV) | OV1288 | MW424401-MW424410 |
